## Supplementary Figures for "Endoplasmic reticulum visits highly active spines and prevents runaway potentiation of synapses" (from another version of this preprint: version 1, 2020-07-30)

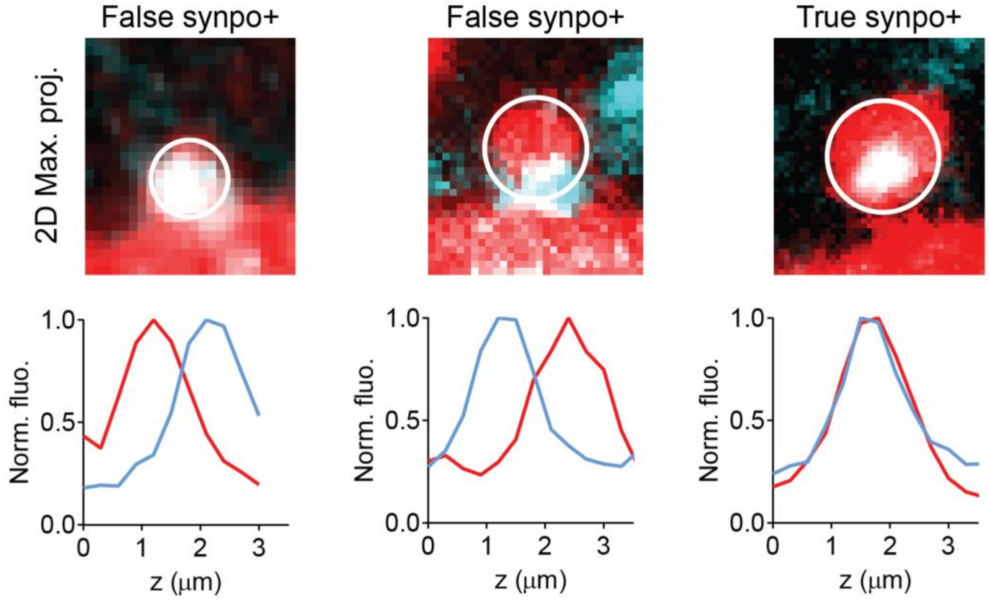


**Supplementary Fig. 1 | Scoring of the spine apparatus marker synaptopodin in fixed tissue.** 2D projections of confocal image z-stacks (upper row) are not sufficient to identify synaptopodin-positive (synpo+) spines. To identify true synpo+ spines, we evaluated the axial intensity profile in confocal image stacks (lower row) in a circular region of interest (white circle). Only spines where an intensity maximum of the anti-synaptopodin signal (cyan) was found in the focal plane of the spine head (red) were scored as synpo+ (right example).

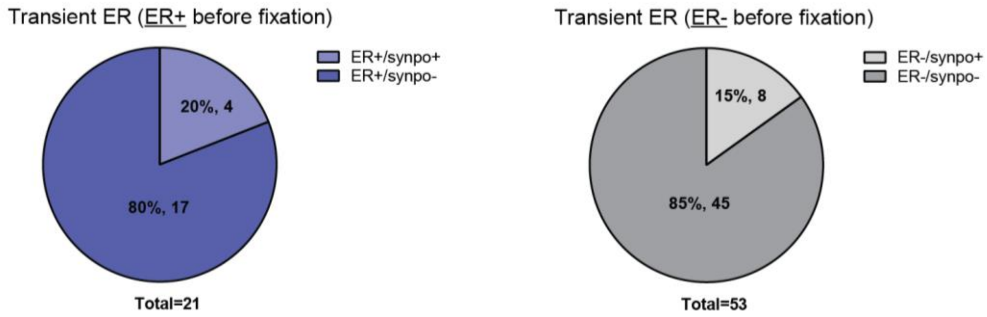


**Supplementary Fig. 2 | Spines with transient ER are rarely immunoreactive for synaptopodin.** To test whether spines that were just invaded by ER were more likely to contain synaptopodin, we split the sample of transient ER spines into the ones that contained ER (ER+, blue) at the last live imaging time point (left diagram) and the ones that did not contain ER (ER-, gray). In both groups, a small percentage was immunoreactive for synaptopodin (20%; 15%).

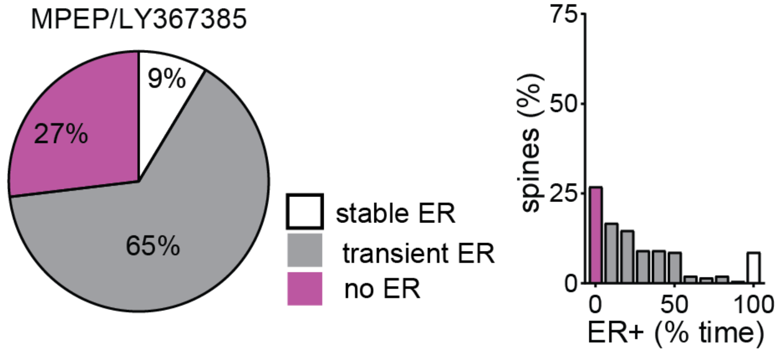


**Supplementary Fig. 3 | Block of group I mGluRs increases ER motility.** Blockade of group I mGluRs with a cocktail of MPEP (50 µM) and LY367385 (100 µM) increased the percentage of spines visited by ER (gray sector) and prolonged visit duration (*n* = 197 spines, 3 cells, 3 slices) compared to controls (see Fig. 1i). Observation time = 100 min.

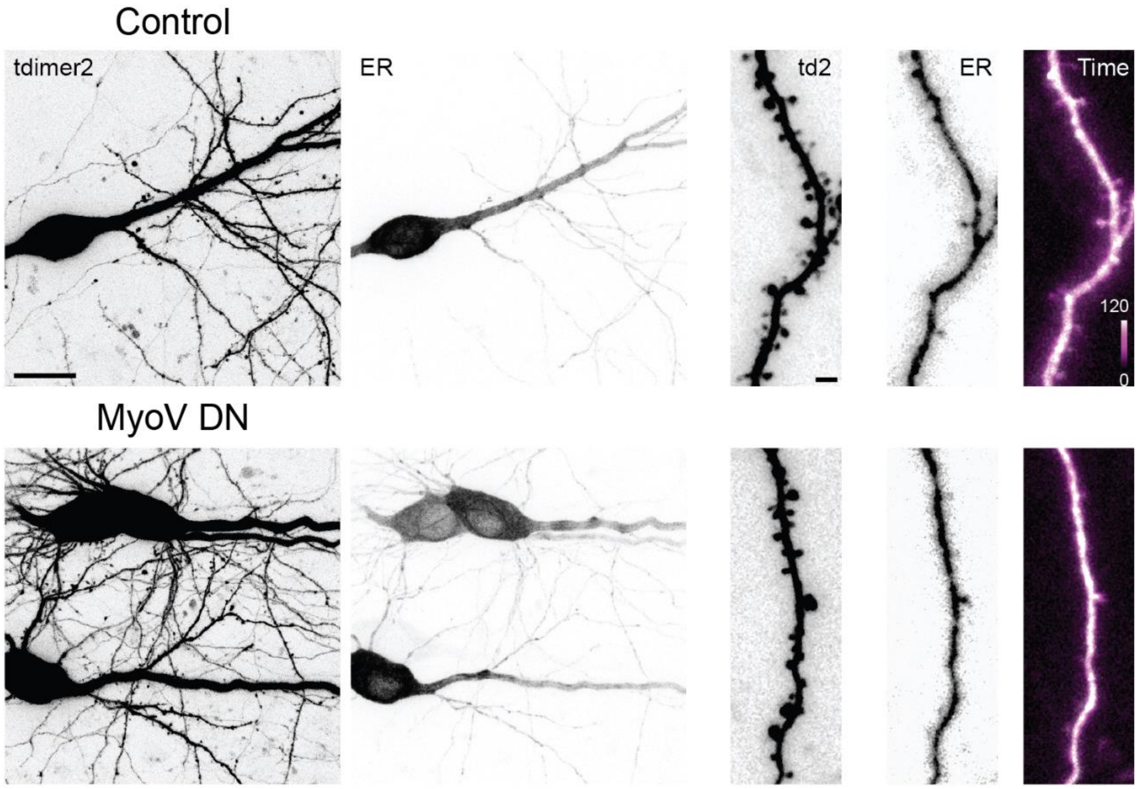


**Supplementary Fig. 4 | Expression of a MyoVa dominant-negative construct in CA1 neurons.**

Maximum projections of two-photon z-stacks of tdimer2 (neuron filler) and ER-EGFP from electroporated CA1 neurons expressing fluorescent reporters (above) or also MyoV DN (below). Scale bar: 20 µm. Detail of dendritic segments (right). Scale bar: 2 µm. Collapsed time projection (far right) of a 120 min time lapse, color coded for the duration of ER signal presence. Duration: 120 min.

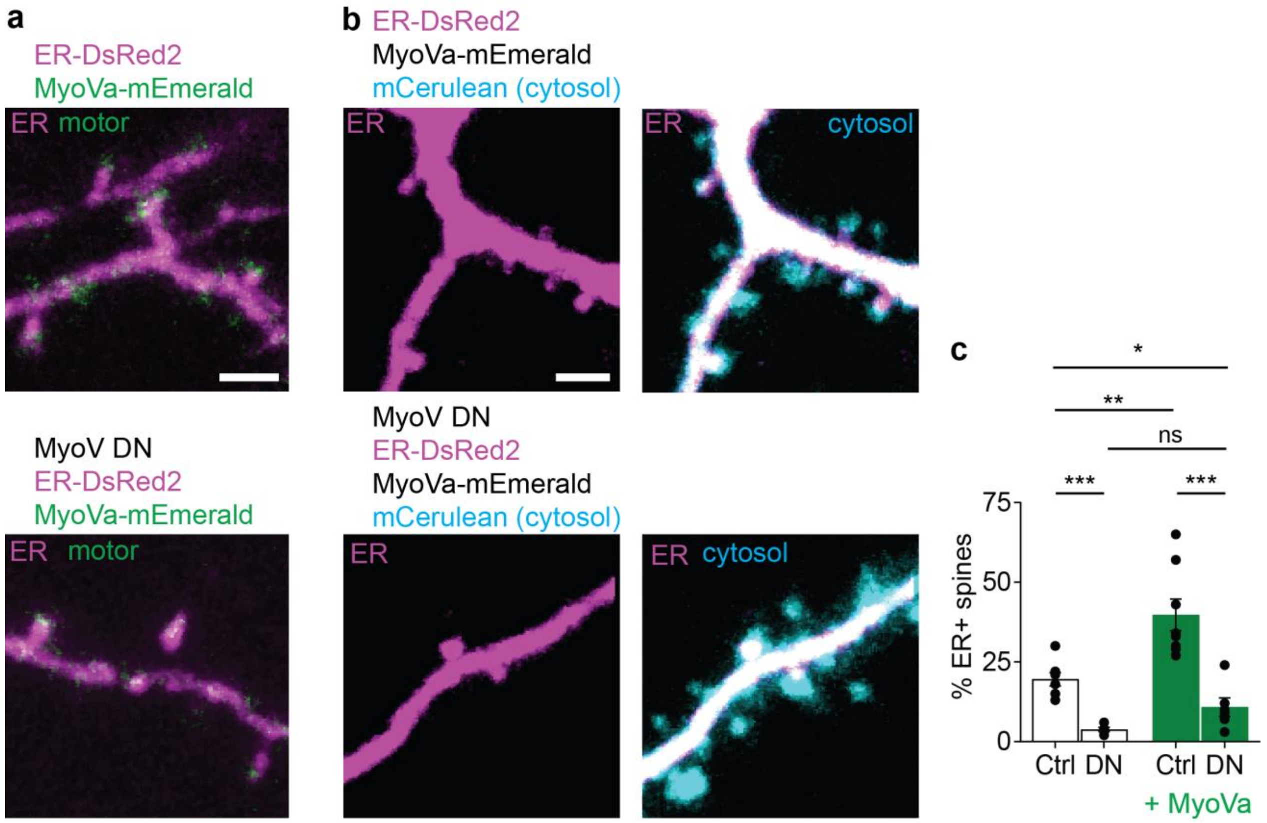


**Supplementary Fig. 5 | Overexpression of MyoVa shows bidirectional modulation of spine ER.**

**a,** Expression of MyoVa-mEmerald shows the localization of the motor at the tip of ER protrusions in dendrites from control (upper image) and MyoV DN-expressing (lower image) CA1 neurons. For these experiments, we labeled the ER with a red fluorescent protein (ER-DsRed2, displayed as magenta). Scale bar: 2 µm. **b,** In neurons overexpressing MyoVa-mEmerald, the dominant negative construct MyoV DN (lower images) still had a strong effect on the number of ER+ spines. Cerulean was used as a cytosolic filler to visualize all spines. Scale bar: 2 µm. **c,** Comparing data (mean ± *SEM*) from Fig. 3c (white bars: control vs MyoV DN, 20 ± 2% vs 4 ± 1% ER+ spines, $p = 0.0001$, unpaired t test) and MyoVa-mEmerald overexpression (green bars: control ($n = 8$ cells, 5 slices) vs MyoV DN ($n = 6$ cells, 4 slices), 40 ± 5% vs 11 ± 3% ER+ spines, $p = 0.0007$). Overexpression of MyoVa-mEmerald doubled the number of ER+ spines under control conditions ($p = 0.004$) and tripled the number of ER+ spines in cells expressing the dominant negative construct ($p = 0.06$). Neurons expressing both MyoVa-mEmerald and the DN construct still have fewer ER+ spines compared to neurons with unperturbed motors (partial rescue, $p = 0.03$).

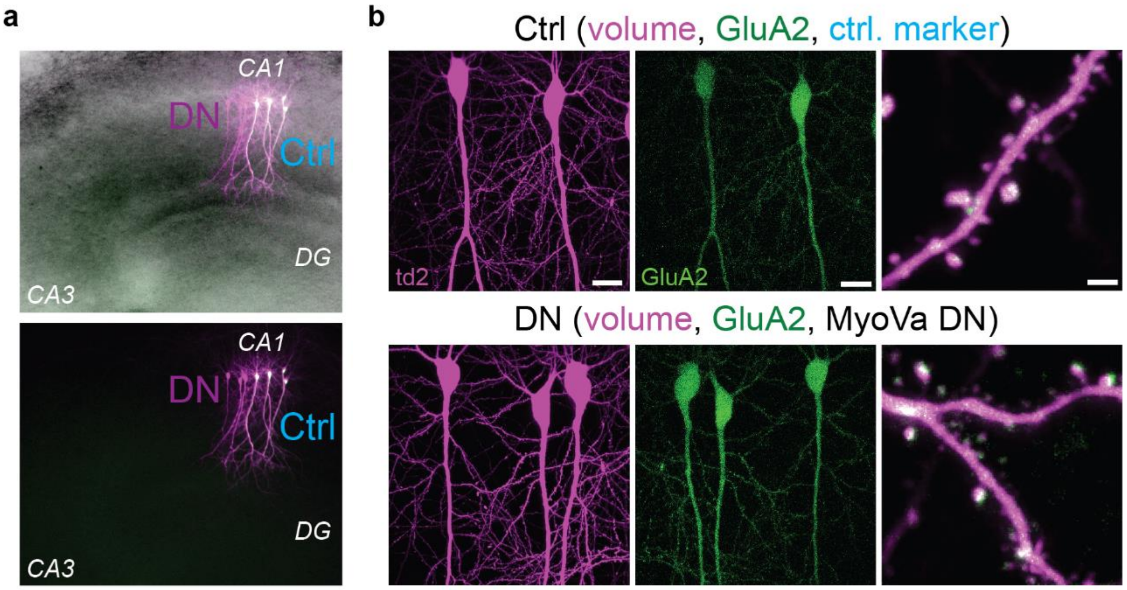


**Supplementary Fig. 6 | Surface expression of AMPA receptors is enhanced in MyoV DN neurons.**

**a,** Overview of organotypic slice culture, showing control neurons expressing SEP-GluA2, cytosolic tdimer2 and cytosolic mCerulean (white), and DN neurons expressing SEP-GluA2, cytosolic tdimer2 and MyoV DN (magenta). **b**, Volume channel (magenta), SEP-labeled AMPA receptors (green) and high magnification overlay (SEP-GluA2 signal appears white). Upper row: control neurons; lower row: MyoV DN neurons. For quantification, see Fig. 4a, b.

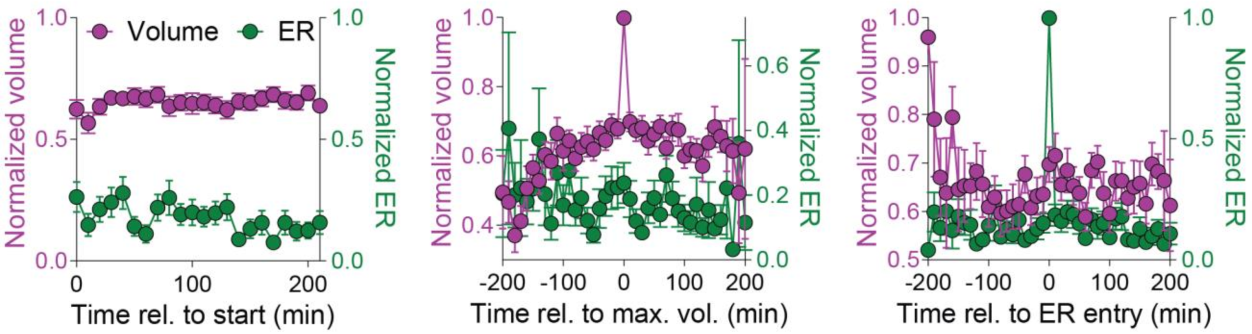


**Supplementary Fig. 7 | ER calcium depletion abolishes correlation between ER entry and spine volume fluctuations.** Thapsigargin (1 µM, 60 min preincubation time), a SERCA pump blocker, preserved ER dynamics and spine volume expansion, but abolished any temporal coupling. *Left,* average of spine volume and ER fluorescence in spines over time. *Center,* at times of maximum spine volume (t = 0), the probability of spines containing an ER was average. *Right,* At the time of ER visit (t = 0), spine volume was average (*n* = 28 spines). These experiments provide evidence that the observed temporal coupling of events in the absence of thapsigargin (Fig. 2e, f) was not an imaging artifact.
